# Genetic and morphological analysis shows the Nechisar Nightjar is a hybrid

**DOI:** 10.1101/2025.04.08.647728

**Authors:** Thomas J. Shannon, Hein van Grouw, J. Martin Collinson

**Affiliations:** School of Medicine, Medical Sciences and Nutrition, University of Aberdeen, Institute of Medical Sciences, Foresterhill, Aberdeen AB25 2ZD

**Keywords:** Caprimulgus, Nechisar Nightjar, genetics, systematics, nightjar

## Abstract

The Nechisar Nightjar *Caprimulgus solala* was described only from a single wing collected as roadkill from Nechisar National Park in Ethiopia in 1990. To resolve its taxonomic status a sample was taken from the Nechisar Nightjar holotype and 53 other Afrotropical nightjars, and genomic DNA extracted for sequencing of one mitochondrial and three nuclear genes. The genetic analysis concluded that Nechisar Nightjar is most likely a hybrid, with mitochondrial DNA of Standard-winged Nightjar *Caprimulgus longipennis* and nuclear alleles from Standard-winged Nightjar and a second nightjar species that has not yet been sequenced. Morphometric and plumage analysis suggests that the paternal parent is Freckled Nightjar *Caprimulgus tristigma*, for which no nuclear sequence data are currently available.

## Introduction

The discovery and description of avian taxa based on single specimens or sets of specimens collected during expeditions is a recurrent theme in ornithology. In many cases, such specimens have been assigned to new species when they do not correspond to known taxa. However, subsequent studies have often shown these unique specimens to be aberrant individuals of more widespread species, or, in some cases, hybrids. For instance, the Liberian Greenbul *Phyllastrephus leucolepis*, initially described from individuals collected in Cavalla Forest, Liberia between 1981 and 1984, was later revealed through genetic investigation to be an individual of the Icterine Greenbul *Phyllastrephus icterinus*, exhibiting an aberrant plumage variation (Collinson et al. 2018). Similarly, Vaurie’s Nightjar *Caprimulgus centralasicus*, described from a single specimen collected in 1929 in Xinjiang, China, was shown to be a female European Nightjar *C. europaeus*, following molecular analyses (Schweizer et al. 2020). These findings underscore the importance of integrating morphological assessment with molecular techniques in resolving the taxonomic identity of such unique specimens.

In some cases, these anomalous specimens may represent hybrid individuals, a possibility that has gained increasing recognition with advances in genetic techniques. A well-documented example involves the Bogotá Sunangel *Heliangelus zusii*, a hummingbird initially described from a skin purchased in Bogotá, Colombia in 1909, and thought to represent a novel taxon. Subsequent genetic analyses, along with the capture of a similar individual in 2011 in Reserva Rogitama, Boyacá, revealed both birds to be hybrids involving the female Long-tailed Sylph *Aglaiocercus kingii* and either a male Tyrian Metaltail *Metallura tyrianthina* (Rogitama individual) or another, as yet genetically unsampled, hummingbird species (Perez-Eman et al. 2018). These cases highlight the potential for hybridization to produce novel phenotypic combinations, which can easily be misinterpreted as new species.

The Nechisar Nightjar *Caprimulgus solala*, described by Safford et al. (1995), represents a similarly enigmatic case. Known solely from a single wing found in 1990 along a dirt road in Nechisar National Park, Ethiopia, this specimen did not match any known species of nightjar. The combination of its wing length and the extent and positioning of a distinct white patch on the outer primaries led to its description as a new species. Given its isolated origin and lack of additional evidence, *C. solala* has been presumed to be a rare and possibly range-restricted species.

Several expeditions to the Nechisar Plains have sought to observe this elusive nightjar in the wild (e.g., Head 2016; Evens et al. 2018), but none have resulted in conclusive sightings. An expedition in 2012, documented by Head (2016), reported a possible encounter with an individual resembling the Nechisar Nightjar, though this observation remains unverified. The continued failure to rediscover the taxon in its presumed habitat, despite repeated efforts in a known conservation area, has led to increasing scepticism regarding the validity of the taxon and calls for further investigation (Forero & Tella 1997).

The question of the Nechisar Nightjar’s status has gained further complexity given the intraspecific variation in plumage that is often observed in Afrotropical nightjars (Jackson 1984). Such variation, which can sometimes exceed interspecific differences, raises the possibility that the Nechisar Nightjar could represent an unusual variant of a more widespread species. Alternatively, the specimen may represent a hybrid individual. Although hybridization has not been documented within Old World nightjars, the nocturnal and cryptic nature of these birds makes the detection of such events particularly challenging. In the absence of additional specimens, molecular analysis of the holotype remains the most viable approach to determining the taxon’s validity.

## Methods

### Sampling

A skin and muscle sample from the Nechisar Nightjar was obtained with the appropriate permits from the Natural History Museum (Tring, UK), enabling the first molecular analysis of this enigmatic species. Given the incomplete sampling of nightjar species across Afrotropical regions, an additional 53 samples from other nightjars were incorporated into the study to provide a broader comparative framework. Feather or blood samples were collected from 39 live specimens, obtained under license by certified bird ringers during routine ringing operations in the UK and abroad. The remaining 14 samples were derived with authorisation from toepads of museum specimens. Collectively, these samples represented 18 of the 26 nightjar species currently recognized by the International Ornithological Congress (IOC) as occurring in Africa.

To enhance the dataset, publicly available sequence data from GenBank were included, ensuring that genetic data for 25 of the 26 African taxa were available for comparison. The one exception was Prigogine’s Nightjar *Caprimulgus prigoginei*, a species known only from a single specimen and whose validity has been questioned (Louette 1990). Although the holotype had previously been sampled, it provided only a limited amount of sequence data (Sonet et al. 2011), and further destructive sampling was deemed inappropriate. A comprehensive list of the samples used in this study is provided in **Table 1**.

**Table 1.** Sample information for all samples used in this study. Genus names: *C.* = *Caprimulgus*; *V.* = *Veles; G.* = *Gactornis*.

| Sample ID | Common Name | Taxon | Accession/ Ring Number | Source |
| --- | --- | --- | --- | --- |
| CAE01 | Egyptian Nightjar | *C. aegyptius aegyptius* | - | This Study |
| CAE02 | Egyptian Nightjar | *C. aegyptius saharae* | - | This Study |
| CCL01 | Long-tailed Nightjar | *C. climacurus* | R: SH10209 | This Study |
| CDO01 | Donaldson-Smith's Nightjar | *C. donaldsoni* | R: AA35401 | This Study |
| CEU01 | European Nightjar | *C. europaeus* | R: O64305 | This Study |
| CEU02 | European Nightjar | *C. europaeus unwini* | R: O35902 | This Study |
| CEU03 | European Nightjar | *C. europaeus europaeus* | - | This Study |
| CEU04 | European Nightjar | *C. europaeus europaeus* | - | This Study |
| CEU05 | European Nightjar | *C. europaeus meridionalis* | - | This Study |
| CEU06 | European Nightjar | *C. europaeus europaeus* | - | This Study |
| CFC01 | Slender-tailed Nightjar | *C. clarus* | M: LIV MUS: T16207 | This Study |
| CFC02 | Slender-tailed Nightjar | *C. clarus* | - | This Study |
| CFC03 | Slender-tailed Nightjar | *C. clarus* | - | This Study |
| CFC04 | Slender-tailed Nightjar | *C. clarus* | - | This Study |
| CFO01 | Square-tailed Nightjar | *C. fossii* | R: AA35409 | This Study |
| CFO02 | Square-tailed Nightjar | *C. fossii* | - | This Study |
| CFR01 | Sombre Nightjar | *C. fraenatus* | R: BB18060 | This Study |
| CFR02 | Sombre Nightjar | *C. fraenatus* | - | This Study |
| CFR03 | Sombre Nightjar | *C. fraenatus* | - | This Study |
| CIN01 | Plain Nightjar | *C. inornatus* | R: AB5254 | This Study |
| CIN02 | Plain Nightjar | *C. inornatus* | R: AB5277 | This Study |
| CLO01 | Standard-winged Nightjar | *C. longipennis* | - | This Study |
| CNA01 | Swamp Nightjar | *C. natalensis* | M: LIV MUS: 12598 | This Study |
| CNG01 | Black-shouldered Nightjar | *C. nigriscapularis* | - | This Study |
| CNG02 | Black-shouldered Nightjar | *C. nigriscapularis* | - | This Study |
| CNJ01 | Nubian Nightjar | *C. nubicus jonesi* | M: NHMUK 1899.8.11.98 | This Study |
| CNN01 | Nubian Nightjar | *C. nubicus nubicus* | M: NHMUK 1904.12.4.5 | This Study |
| CNR01 | Nubian Nightjar | *C. nubicus torridus* | M: NHMUK 1969.46.20 | This Study |
| CNR02 | Nubian Nightjar | *C. nubicus torridus* | M: NHMUK 1947.64.4 | This Study |
| CNT01 | Nubian Nightjar | *C. nubicus tamaricis* | - | This Study |
| CNT07 | Nubian Nightjar | *C. nubicus tamaricis* | M: NHMUK 1923.8.7.4973 | This Study |
| CNT08 | Nubian Nightjar | *C. nubicus tamaricis* | M: NHMUK 1977.1.4 | This Study |
| CRD01 | Red-necked Nightjar | *C. ruficollis desertorum* | M: NHMUK 1911.12.23.888 | This Study |
| CRR30 | Red-necked Nightjar | *C. ruficollis ruficollis* | - | This Study |
| CRR31 | Red-necked Nightjar | *C. ruficollis ruficollis* | - | This Study |
| CRU01 | Rwenzori Nightjar | *C. poliocephalus ruwenzorii* | R: A2-043 | This Study |
| CRU02 | Rwenzori Nightjar | *C. poliocephalus ruwenzorii* | R: A3-054 | This Study |
| CSO01 | Nechisar Nightjar | *C. solala* | M: NHMUK 1992.6.1 | This Study |
| **CST01** | Star-spotted Nightjar | *C. stellatus* | M: NHMUK 1991.12.2 | This Study |
| **CST02** | Star-spotted Nightjar | *C. stellatus* | - | This Study |
| **CST03** | Star-spotted Nightjar | *C. stellatus* | - | This Study |
| **CST04** | Star-spotted Nightjar | *C. stellatus* | - | This Study |
| **CTR03** | Freckled Nightjar | *C. tristigma* | M: NHMUK 1945.34.750 | This Study |
| **CV01** | European Nightjar | *C. europaeus* | R: 71009192 | This Study |
| **CV02** | European Nightjar | *C. europaeus* | R: 91009211 | This Study |
| **CV03** | European Nightjar | *C. europaeus* | R: 91009212 | This Study |
| **CV04** | European Nightjar | *C. europaeus* | R: 71009232 | This Study |
| **CV05** | European Nightjar | *C. europaeus* | - | This Study |
| **CV06** | European Nightjar | *C. europaeus* | R: 71009247 | This Study |
| **CV07** | European Nightjar | *C. europaeus* | R: 71009248 | This Study |
| **CV08** | European Nightjar | *C. europaeus* | - | This Study |
| **VB01** | Brown Nightjar | *V. binotatus* | M: NHMUK 1977.20.293 | This Study |
| **GU586631** | Egyptian Nightjar | *C. aegyptius* | M: LSUMZ B46332 | Genbank |
| **GU586636** | Bates' Nightjar | *C. batesi* | M: USNM B9899 | Genbank |
| **GU586639** | Square-tailed Nightjar | *C. fossii* | M: ZMUC 115493 | Genbank |
| **DQ062144** | Long-tailed Nightjar | *C. climacurus* | - | Genbank |
| **GU586687** | Long-tailed Nightjar | *C. climacurus* | M: FMNH 396431 | Genbank |
| **GU586654** | Rufous-cheeked Nightjar | *C. rufigena* | M: MBM 7950 | Genbank |
| **DQ062136** | Sombre Nightjar | *C. fraenatus* | - | Genbank |
| **GU586674** | Pennant-winged Nightjar | *C. vexillarius* | - | Genbank |
| **DQ062143** | Standard-winged Nightjar | *C. longipennis* | - | Genbank |
| **GU586673** | Standard-winged Nightjar | *C. longipennis* | M: KUNHM 15546 | Genbank |
| **DQ062138** | Plain Nightjar | *C. inornatus* | - | Genbank |
| **AJ004074** | European Nightjar | *C. europaeus* | - | Genbank |
| **DQ062139** | European Nightjar | *C. europaeus* | - | Genbank |
| **DQ062147** | European Nightjar | *C. europaeus* | - | Genbank |
| **DQ062135** | European Nightjar | *C. europaeus* | - | Genbank |
| **GU586650** | Fiery-necked Nightjar | *C. pectoralis* | M: UWBM 71315 | Genbank |
| **GU586651** | Montane Nightjar | *C. poliocephalus poliocephalus* | M: MBM 11252 | Genbank |
| **GU586647** | Black-shouldered Nightjar | *C. nigriscapularis* | M: AMNH DOT 10638 | Genbank |
| **GU586646** | Madagascar Nightjar | *C. madagascariensis* | M: FMNH 436426 | Genbank |
| **LT671509** | Golden Nightjar | *C. eximius* | - | Genbank |
| **GU586637** | Collared nightjar | *G. enarratus* | M: FMNH 431158 | Genbank |

### Molecular Techniques

Total genomic DNA was isolated from 1-3 feather tips, 1-2 mm³ of blood, or 1-2 mm³ of tissue per individual using the QIAamp DNA Micro Kit (QIAGEN, UK) as per manufacturer’s instructions, with the addition of dithiothreitol to 0.1 M concentration into the digestion mix and elution in 80 μl of QIAGEN Buffer AE.

A single mitochondrial locus, cytochrome b (*cytb*), was amplified by PCR using primers and protocols as in Helbig et al. (1995) for fresh samples. For degraded museum specimens, a series of overlapping short fragment primer sets were designed based on consensus sequence from all *Caprimulgus* sp. data available in GenBank. These primer pairs were also amplified using the protocol outlined in Helbig & Seibold (1999). Three nuclear loci were also targeted: MYC, RP1L1 and REST. A full list of loci used in this study is shown in **Table 2**. Primers and protocols described in Han et al. (2010) for MYC and Liu et al. (2018) for RP1L1 and REST were followed, with a bespoke forward primer designed for the REST gene based on existing Caprimulgid sequence data available on Genbank. A full list of primers used is provided in **Table 3**. PCR products were gel purified using the QIAquick Gel Extraction Kit (QIAGEN, UK), before being sent for Sanger sequencing at Source Bioscience (Nottingham, UK).

**Table 2.** Genes targeted in this study, along with primers and PCR protocols followed.

| Locus | Gene Name | Approximate Length (bp) | Type |
| --- | --- | --- | --- |
| **Cytb** | Cytochrome b | 1050 | Mitochondrial |
| **CHD1-Z/W** | Chromo-helicase-DNA binding 1 | 450-600 | Sex-linked |
| **REST** | RE1-Silencing Transcription gene | 600 | Autosomal Nuclear |
| **RP1L1** | Retinitis pigmentosa 1 like 1 | 600 | Autosomal Nuclear |
| **MYC** | Cellular myelocytomatosis oncogene | 400 | Autosomal Nuclear |

**Table 3.**
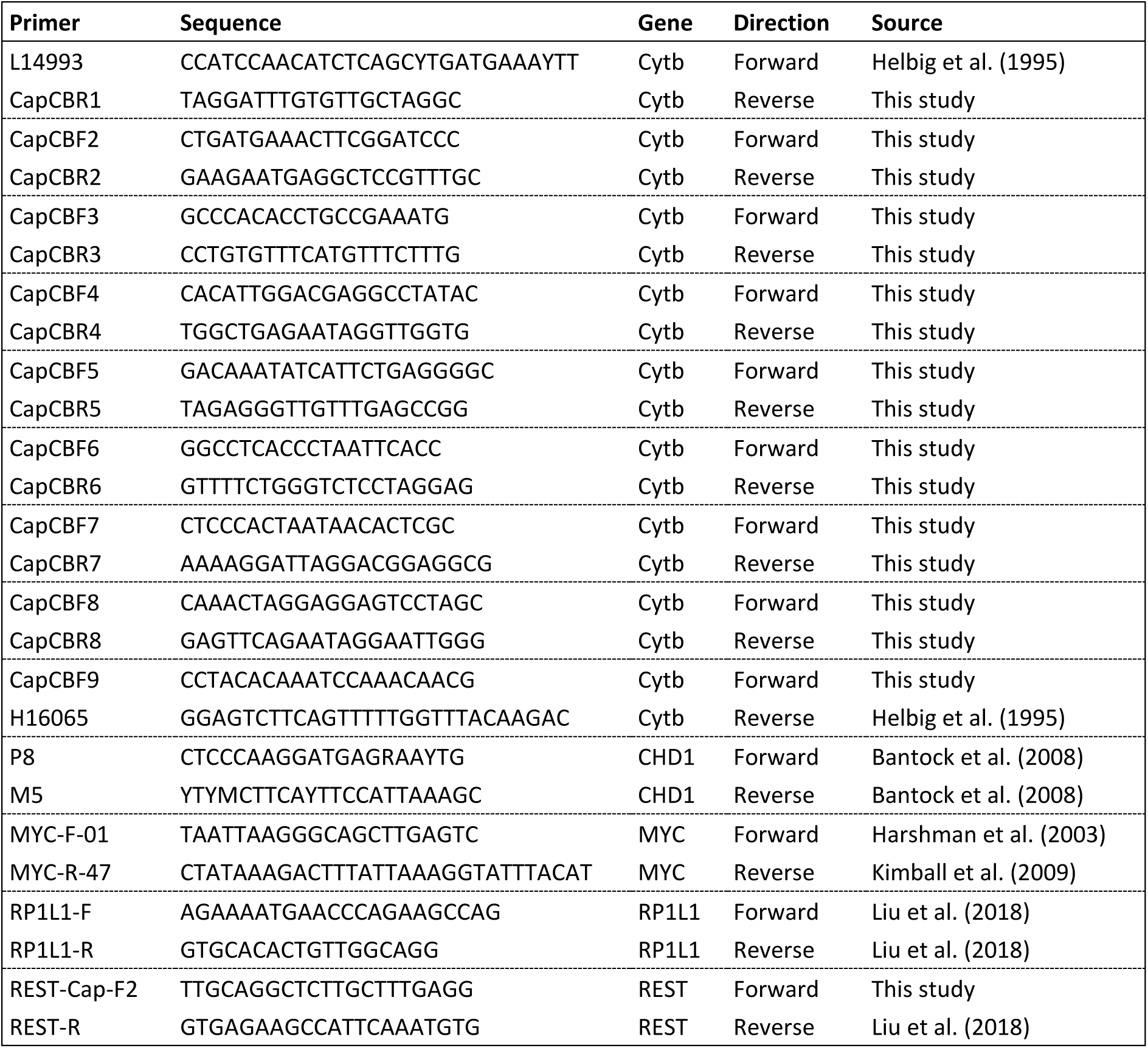
A complete list of primers used in this study.

Molecular sexing of the Nechisar Nightjar holotype was conducted using the 2550F/2718R primer method targeting the CHD1 gene for non-ratite birds, as outlined in Fridolfsson & Ellegren (1999).

Extreme precautions were taken to avoid cross contamination throughout. The Nechisar Nightjar sample was extracted and sequenced before any other nightjar material, and internal muscle tissue was used to eliminate the remote possibility of surface DNA contamination. Fresh samples and museum specimens were processed separately, blank extractions performed, and clean UV-irradiated hoods and plastics were used throughout. No museum specimen ever produced a PCR band larger than 220 bp that would be indicative of contamination with modern DNA.

All verified sequences have been uploaded to the European Nucleotide Archive, Accession numbers tba upon final publication.

### Phylogenetic analyses

All Sanger sequences were checked by eye, and for museum samples, overlapping short fragment sequences were concatenated in CLC Genomics Workbench 12 (QIAGEN, UK) to give a single consensus cytb sequence with all primers trimmed out. Cytochrome b sequences were aligned in CLC Genomics Workbench 12 (QIAGEN, UK) and substitution models assessed in MEGA X (Kumar et al. 2018). The best model was selected based on the Bayesian Information Criterion (BIC).

A phylogeny was produced with BEAST v. 2.6.1 (Bouckaert et al. 2014). The TN93+G model was selected, with five gamma rate categories. The analysis utilised default priors and operators, followed a yule model and was run for 10 million generations, sampling every 1000 generations.

The output of the run was inspected in Tracer v. 1.7.1 (Rambaut et al. 2018) to ensure all effective samples sizes (ESS) were above 200. A 10% burn-in was selected and a maximum clade credibility tree produced using median node heights in TreeAnnotator (Drummond & Rambaut 2007). This tree was then visualised in FigTree 1.4.4 (Rambaut 2018).

To investigate consistency between tree-building methods, a Maximum Likelihood phylogeny was produced in MEGA X (Kumar et al. 2018), using the same model parameters as above. A total of 100 bootstrap replicates were run to assess the statistical confidence of the final tree.

Nuclear sequences were checked by eye, and haplotypes from the Nechisar Nightjar were separated manually. Samples were then aligned and similarity of each Nechisar haplotype compared against sequences from all other Afrotropical nightjar species to identify parentage. Sequences were uploaded to Genbank.

### Morphometrics

Mean wing lengths, white patch positions and P9 emargination positions of Nechisar Nightjar and other Afrotropical Nightjars were taken from Safford et al. (1995) and Jackson (2000).

## Results

### Mitochondrial DNA

In an initial analysis, cytb sequence was obtained from the Nechisar Nightjar sample and compared to that of all other available nightjar species. The Nechisar Nightjar holotype had mitochondrial DNA that showed uncorrected 99.42-99.54% identity for the only two Standard-winged Nightjar *C. longipennis* cytb sequences in Genbank, accession numbers GU586673 (1026/1031 bp match) and DQ062143 (650/653 bp match). All other available nightjars were very (6-13%) divergent.

In light of this result, and given that sequences from some nightjar taxa were not publicly available, further Standard-winged Nightjar other *Caprimulgus* specimens were obtained for phylogenetic analysis as described in the Methods Section. The Maximum Likelihood and BEAST phylogenies (Clade A in **Figure 1** & Supplementary **Figure 1**) placed the holotype confidently within Standard-winged Nightjar clade (99% bootstrap/posterior probability of 1. The specimen’s maternal lineage evidently comes from Standard-winged Nightjar, indicating that the holotype is either a Standard-winged Nightjar with aberrant plumage or a hybrid.

**Figure 1.**
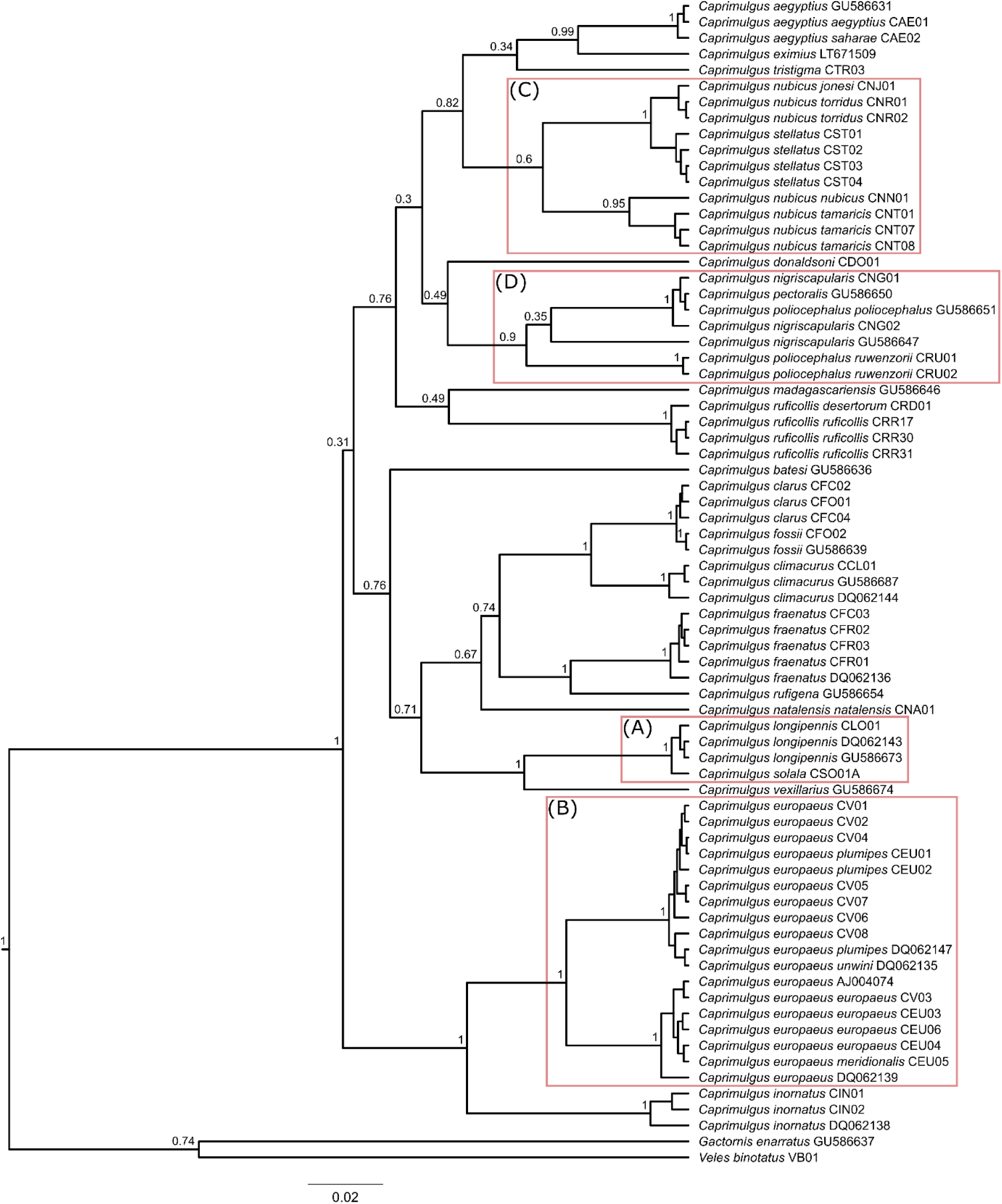
A phylogeny of European and Afrotropical nightjars reconstructed in BEAST based on the mitochondrial cytochrome b gene. Numbers at nodes represent posterior probabilities. Clades of note are highlighted in red boxes: (A) Nechisar Nightjar is, at least maternally, a Standard-winged Nightjar; (B) deep divergence observed within European Nightjar; (C) paraphyly in Nubian Nightjar complex; (D) poor resolution in the Black-shouldered/Fiery-necked/Montane Nightjar complex

Beyond the placement of Nechisar Nightjar, there were a number of other notable findings, including the deep mitochondrial lineage divergenge within European Nightjar separating eastern and western populations (Clade B in **Figure 1** & Supplementary **Figure 1**), and the species’ position as sister to Plain Nightjar *C. inornatus*. Perhaps most unexpected was the position of Star-spotted Nightjar *C. stellatus* within the Nubian Nightjar *C. nubicus* clade (Clade C in **Figure 1** & Supplementary **Figure 1**). This suggests that there may be paraphyly within Nubian Nightjar, with the pale migratory northern populations from Sudan and the Levant representing a separate taxon from the darker sedentary populations of East Africa and Socotra. However posterior probabilities/bootstrap values for higher nodes within this clade show lower levels of support, indicating further genomic work is required to more fully understand the evolutionary history of these taxa. Additionally, there is a lack of logical structure in the Black-shouldered/Fiery-necked/Montane Nightjar (*C. nigriscapularis/C. pectoralis/C. poliocephalus*) complex (Clade D in **Figure 1 Supplementary Figure 1**). This complex also warrants further study to generate a more robust phylogeny

#### 5.3.2 Nuclear DNA

Datasets were collected for three nuclear loci (see **Table 4** for details). Nuclear sequence from at least one locus was acquired for 22 of the 26 European and Afrotropical Nightjar species. Prigogine’s Nightjar holotype was not included in this study, while the samples for Brown Nightjar *Veles binotatus*, Freckled Nightjar *C. tristigma* and Swamp Nightjar *C. natalensis* were old museum specimens which failed to yield any nuclear sequence. For successful samples, 728 bp of sequence was obtained for MYC, 772 bp for RP1L1, and 542 bp for REST. Nechisar Nightjar sequence was resolved as heterozygous between two haplotypes, one of which (Haplotype 1) was consistent with Standard-winged Nightjar and the other (Haplotype 2) was ‘unknown’ and compared to other nightjar taxa.

**Table 4.** Species list of all Eurasian and Afrotropical nightjar species, detailing for which taxa nuclear and mitochondrial sequence data was acquired for this study. No nuclear material was recovered for four species, and these are therefore not included in the genetic analysis of the parentage of Nechisar Nightjar. A single species, Prigogine’s Nightjar, known from a single specimen could not be sampled, and is therefore also excluded from the wider phylogeny.

| Common Name | Species Name | MYC | RP1L1 | REST | CYTB |
| --- | --- | --- | --- | --- | --- |
| Brown Nightjar | *V. binotatus* | - | - | - | ✓ |
| Red-necked Nightjar | *C. ruficollis* | ✓ | ✓ | ✓ | ✓ |
| European Nightjar | *C. europaeus* | ✓ | ✓ | ✓ | ✓ |
| Sombre Nightjar | *C. fraenatus* | ✓ | ✓ | ✓ | ✓ |
| Rufous-cheeked Nightjar | *C. rufigena* | ✓ | - | - | ✓ |
| Egyptian Nightjar | *C. aegyptius* | ✓ | ✓ | ✓ | ✓ |
| Nubian Nightjar | *C. nubicus* | - | - | ✓ | ✓ |
| Golden Nightjar | *C. eximius* | ✓ | - | - | ✓ |
| Donaldson Smith's Nightjar | *C. donaldsoni* | ✓ | ✓ | ✓ | ✓ |
| Black-shouldered Nightjar | *C. nigriscapularis* | ✓ | ✓ | ✓ | ✓ |
| Fiery-necked Nightjar | *C. pectoralis* | ✓ | - | - | ✓ |
| Montane Nightjar | *C. poliocephalus* | ✓ | ✓ | ✓ | ✓ |
| Madagascar Nightjar | *C. madagascariensis* | ✓ | - | - | ✓ |
| Swamp Nightjar | *C. natalensis* | - | - | - | ✓ |
| Nechisar Nightjar | *C. solala* | ✓ | ✓ | ✓ | ✓ |
| Plain Nightjar | *C. inornatus* | - | ✓ | ✓ | ✓ |
| Star-spotted Nightjar | *C. stellatus* | - | ✓ | ✓ | ✓ |
| Freckled Nightjar | *C. tristigma* | - | - | - | ✓ |
| Prigogine's Nightjar | *C. prigoginei* | - | - | - | - |
| Bates's Nightjar | *C. batesi* | ✓ | - | - | ✓ |
| Long-tailed Nightjar | *C. climacurus* | ✓ | ✓ | ✓ | ✓ |
| Slender-tailed Nightjar | *C. clarus* | - | ✓ | ✓ | ✓ |
| Square-tailed Nightjar | *C. fossii* | ✓ | ✓ | ✓ | ✓ |
| Standard-winged Nightjar | *C. longipennis* | ✓ | ✓ | ✓ | ✓ |
| Pennant-winged Nightjar | *C. vexillarius* | ✓ | - | - | ✓ |
| Collared Nightjar | *G. enarratus* | ✓ | - | - | ✓ |

**Table 5** details the top 5 hits from the dataset for each haplotype for all three nuclear loci based on percentage similarity. All nightjars are very similar across these short fragments of nuclear loci. Nevertheless, Haplotype 1 was consistently a best match for Standard-winged Nightjar across all 3 nuclear loci, with 99.27-99.82% identity. For all 3 loci, Haplotype 2 had a lower percentage identity to the top hit than Haplotype 1. There was also no consistency in the top matching taxon for Haplotype 2 (*C. rufigena*, *C. fraenatus* and *C. clarus* for MYC, RP1L1 and REST, respectively). This indicates that Nechisar Nightjar’s paternal taxon is likely to be absent from the nuclear dataset.

**Table 5.** BLAST search results for each Nechisar Nightjar haplotype for the three nuclear loci. Haplotype 1 refers to the Standard-winged Nightjar haplotype in all loci, while haplotype 2 comes from the paternal lineage. Only the top 5 hits in terms of similarity are shown.

| MYC 728 bp |  |  |  |
| --- | --- | --- | --- |
| *C. solala* Haplotype 1 |  | *C. solala* Haplotype 2 |  |
| Taxon | % identity | Taxon | % identity |
| *C. longipennis* | 99.73 | *C. rufigena* | 99.20 |
| *C. vexillarius* | 99.47 | *C. longipennis* | 99.20 |
| *C. rufigena* | 98.93 | *C. vexillarius* | 98.93 |
| *C. pectoralis* | 98.93 | *C. fossii* | 98.93 |
| *C. fossii* | 98.66 | *C. pectoralis* | 98.40 |
| RP1L1 772 bp |  |  |  |
| *C. solala* Haplotype 1 |  | *C. solala* Haplotype 2 |  |
| Taxon | % identity | Taxon | % identity |
| *C. longipennis* | 99.27 | *C. fraenatus* | 99.13 |
| *C. aegyptius* | 98.69 | *C. aegyptius* | 99.13 |
| *C. poliocephalus* | 98.40 | *C. poliocephalus* | 98.98 |
| *C. climacurus* | 98.26 | *C. climacurus* | 98.98 |
| *C. inornatus* | 98.26 | *C. inornatus* | 98.84 |
| REST 542 bp |  |  |  |
| *C. solala* Haplotype 1 |  | *C. solala* Haplotype 2 |  |
| Taxon | % identity | Taxon | % identity |
| *C. longipennis* | 99.82 | *C. clarus* | 99.63 |
| *C. nigriscapularis* | 99.26 | *C. fraenatus* | 99.45 |
| *C. donaldsoni* | 99.26 | *C. nigriscapularis* | 99.45 |
| *C. ruficollis* | 99.08 | *C. donaldsoni* | 99.45 |
| *C. fraenatus* | 99.08 | *C. ruficollis* | 99.26 |

**Table 6.** Range and morphometric similarity of male nightjars to the Nechisar Nightjar holotype. The holotype has white patches on 4 outer primaries, and on 2 outer tail feathers. Wing lengths within 10% of the Nechisar holotype (188mm) were considered a match. Matching criteria shown in bold, taxa which match all criteria are highlighted in green. Wing lengths (longest primary measurement) are from Jackson (2000) and ranges from Holyoak and Woodcock (2001),

| Common name | Species name | Occurs in Ethiopia | # Primaries with white patch | White patch on 2 outer tail feathers | Wing length (mm) | WL range (mm) |
| --- | --- | --- | --- | --- | --- | --- |
| Freckled Nightjar | *C. tristigma* | Y | 4 | Y | **184.2** | **168-187** |
| Sombre Nightjar | *C. fraenatus* | Y | 4 | Y | 163.2 | 152-172 |
| Fiery-necked Nightjar | *C. pectoralis* | Y | 4 | Y | 162 | 152-177 |
| Plain Nightjar | *C. inornatus* | Y | 4 | Y | 159.8 | 152-164 |
| Star-spotted Nightjar | *C. stellatus* | Y | 4 | Y | 154.9 | 151-162 |
| Swamp Nightjar | *C. natalensis* | Y | 4 | Y | 151.5 | 148-162 |
| Nubian Nightjar | *C. nubicus* | Y | 4 | Y | 151.4 | 141-157 |
| Donaldson Smith's Nightjar | *C. donaldsoni* | Y | 4 | Y | 132.5 | 119-145 |
| Egyptian Nightjar | *C. aegyptius* | Y | 0 | Y | **196** | **193-216** |
| European Nightjar | *C. europaeus* | Y | 3 | Y | **190** | **175-201** |
| Montane Nightjar | *C. poliocephalus* | Y | 4 | N | 152.3 | 144-163 |
| Standard-winged Nightjar | *C. longipennis* | Y | 0 | N | **173.9** | **166-181** |
| Slender-tailed Nightjar | *C. clarus* | Y | 6 | N | 147.1 | 133-155 |
| Long-tailed Nightjar | *C. climacurus* | Y | 5 | N | 143.2 | 129-151 |
| Pennant-winged Nightjar | *C. vexillarius* | N | 8 | N | 221.2 | 200-266 |
| Red-necked Nightjar | *C. ruficollis* | N | 3 | Y | **201.1** | **198-214** |
| Bates's Nightjar | *C. batesi* | N | 4 | Y | **189.3** | **180-201** |
| Brown Nightjar | *V. binotatus* | N | 0 | Y | 151.1 | 140-163 |
| Golden Nightjar | *C. eximius* | N | 4 | Y | **175.1** | **169-186** |
| Prigogine's Nightjar | *C. prigoginei* | N | 3 | Y | 164 | (164) |
| Rufous-cheeked Nightjar | *C. rufigena* | N | 4 | Y | 162.3 | 154-169 |
| Square-tailed Nightjar | *C. fossii* | N | 5 | N | 154 | 161-168 |
| Madagascar Nightjar | *C. madagascariensis* | N | 4 | Y | 153.7 | 147-163 |
| Black-shouldered Nightjar | *C. nigriscapularis* | N | 4 | Y | 151.8 | 147-159 |
| Collared Nightjar | *G. enarratus* | N | 0 | N | 148.6 | 147-152 |

### Morphometrics

The wing of Nechisar Nightjar was rexamined.

The Nechisar Nightjar holotype has white patches on the four outermost primaries, strongly suffused with buff and cinnamon tones, and was noted as having white patches on at least the 2 outermost tail feathers though these were not collected along with the wing from which the species was described (Safford et al. 1995). Based on the genetic data, the Nechisar holotype is a hybrid between a female Standard-winged Nightjar and another nightjar species for which no nuclear sequence has yet be published or obtained by us. Following this conclusion, with the lack of a plausible genetic candidate for the paternal taxon, a morphological study was conducted to deduce the most likely father. The molecular sexing showed the holotype to be a male, consistent with the presence of white patches in the wings and tail, so morphological characters exhibited by males of each species were considered. Hybrids often are not ‘simple’ intermediate between their parent species, but with this caveat in mind, the following analysis was undertaken.

Of the Eurasian and Afrotropical nightjar species, 14 (including Standard-winged Nightjar) are known to occur in Ethiopia. Of those 14 species, nine have a white patch on exactly four of the outermost primary feathers. Eight of those nine species also have white patches on at least the two outermost tail feathers. Seven of the remaining eight species have wing lengths much smaller than the Nechisar holotype (132.5-163.2 mm, relative to a wing length of 188 mm for the holotype). While it is not uncommon for hybrids to be larger than both parent species, this left only a single species which matched all morphological criteria, and had an average wing length within 10% of the Nechisar holotype: Freckled Nightjar. This species was also absent from the nuclear DNA study. Additional Freckled Nightjar samples had been procured from Liverpool World Museum, UK, and the National Museum, Bloemfontein, South Africa, for this study, as the NHM specimen we had sampled had produced a low-quality DNA yield. However, the Liverpool World Museum sample failed to yield any DNA, as their museum has a history of arsenic use for skin preservation, while the specimen sampled from the National Museum, Bloemfontein, produced a good quality yield, but was found to be an erroneously labelled European Nightjar.

To further test the hypothesis that the parentage of the Nechisar holotype was Freckled Nightjar x Standard-winged Nightjar *C. tristigma x longipennis*, the position of the emargination on the second outermost primary feather (P9) was investigated in these two species and the Nechisar holotype. This feature has been shown to be important in the identification of nightjars (Jackson, 2000). The ratio of the average distance of the emargination of P9 from the carpal relative to the average distance of the emargination to the feather tip was calculated for each taxon and then used to identify the expected emargination position on a P9 feather matching the length seen in the Nechisar holotype. In **Figure 3**, it is shown that the emargination position of the Nechisar holotype is almost exactly equidistant between the expected emargination positions of Standard-winged and Freckled Nightjars of an equal size as the Nechisar holotype.

**Figure 2:**
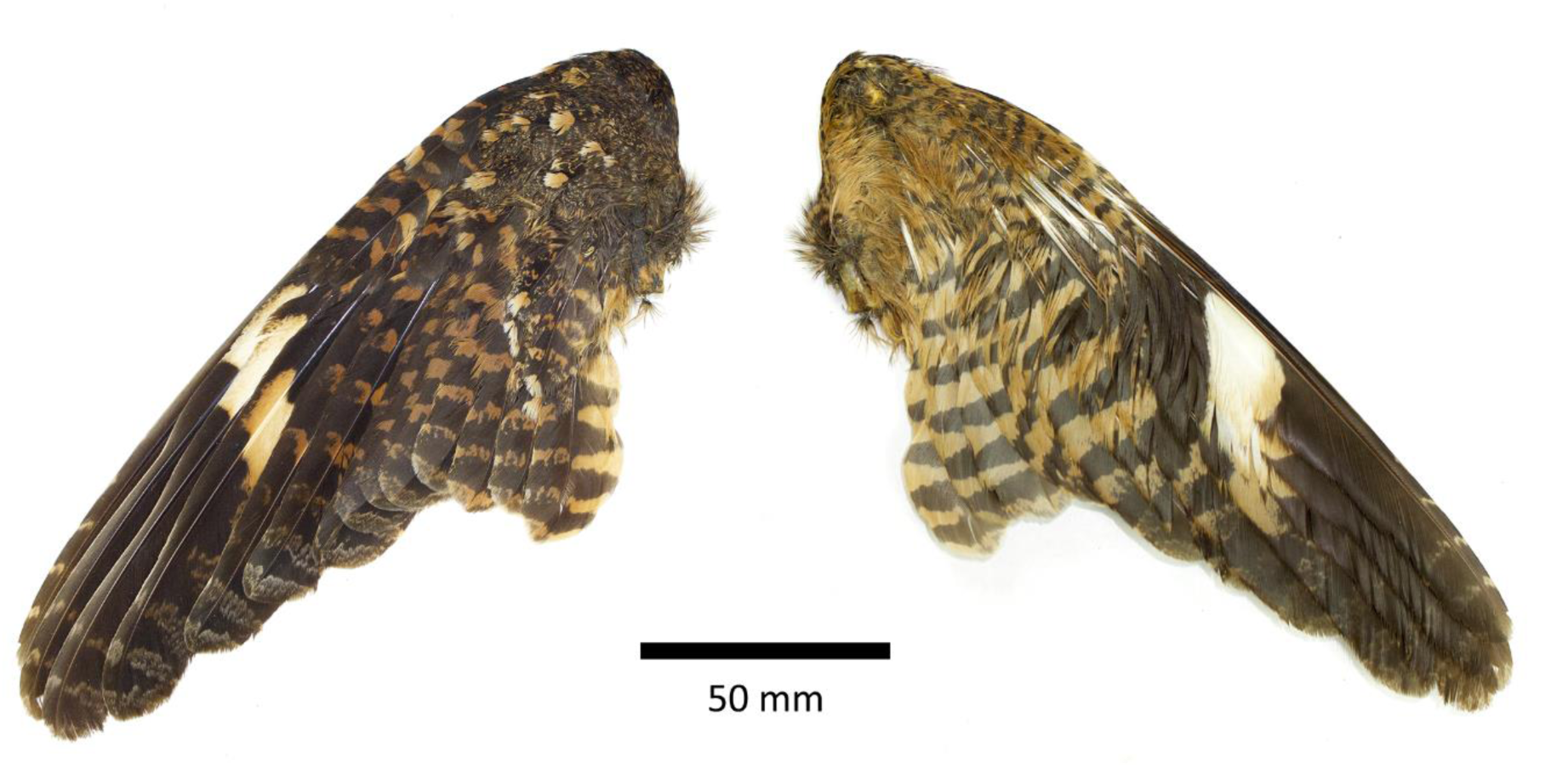
Montage of the upper (left) and underwing (right) of the Nechisar Nightjar holotype.

**Figure 3.**
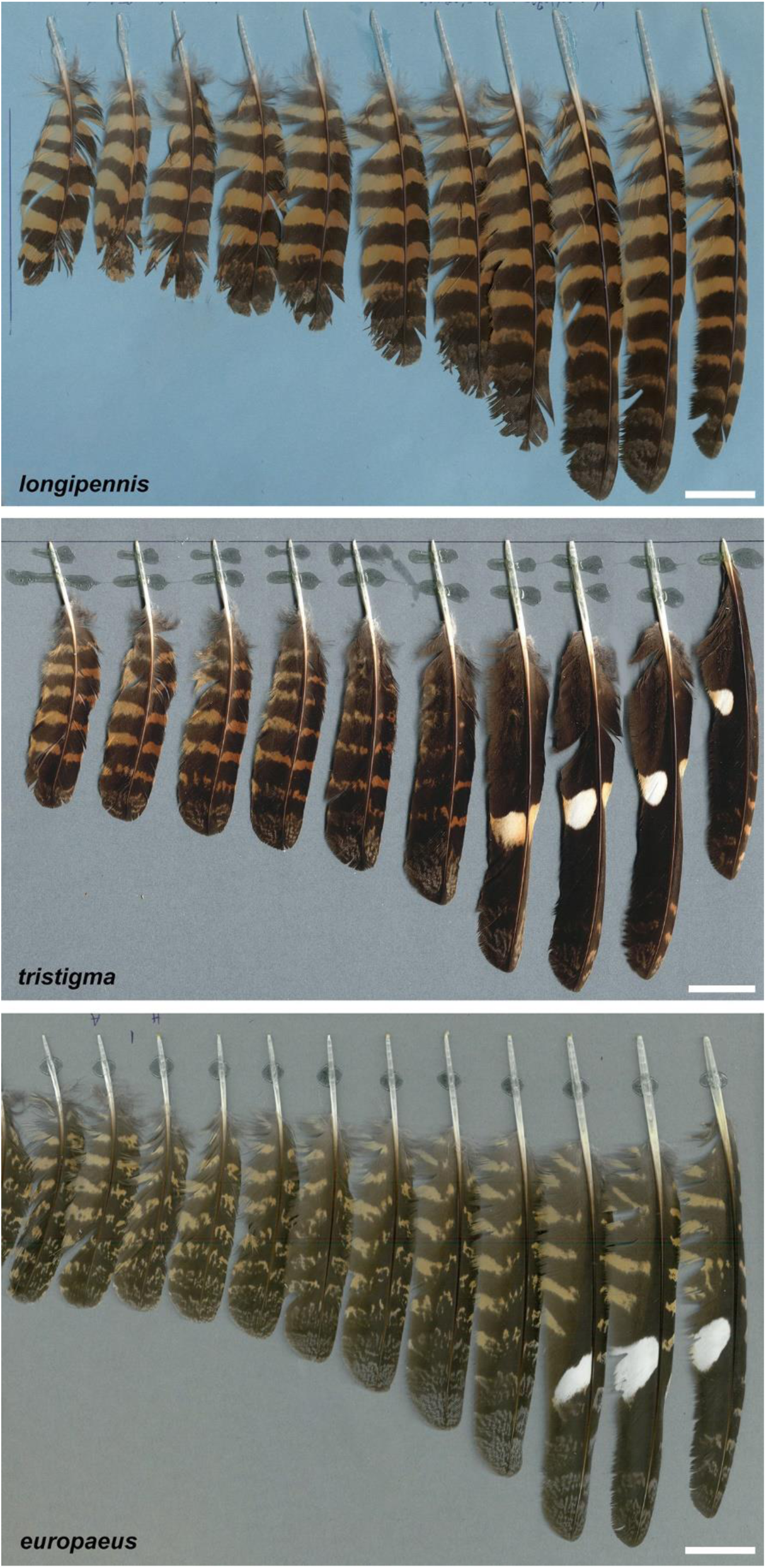
Primary feathers of (from top) representative Standard-winged Nightjars, Freckled Nightjar and European Nightjar. Modified from images uploaded at *Featherbase* (https://www.featherbase.info/en/home), where copyright is retained by the publishers and authors. Scale bars represent 20 mm.

**Figure 4.**
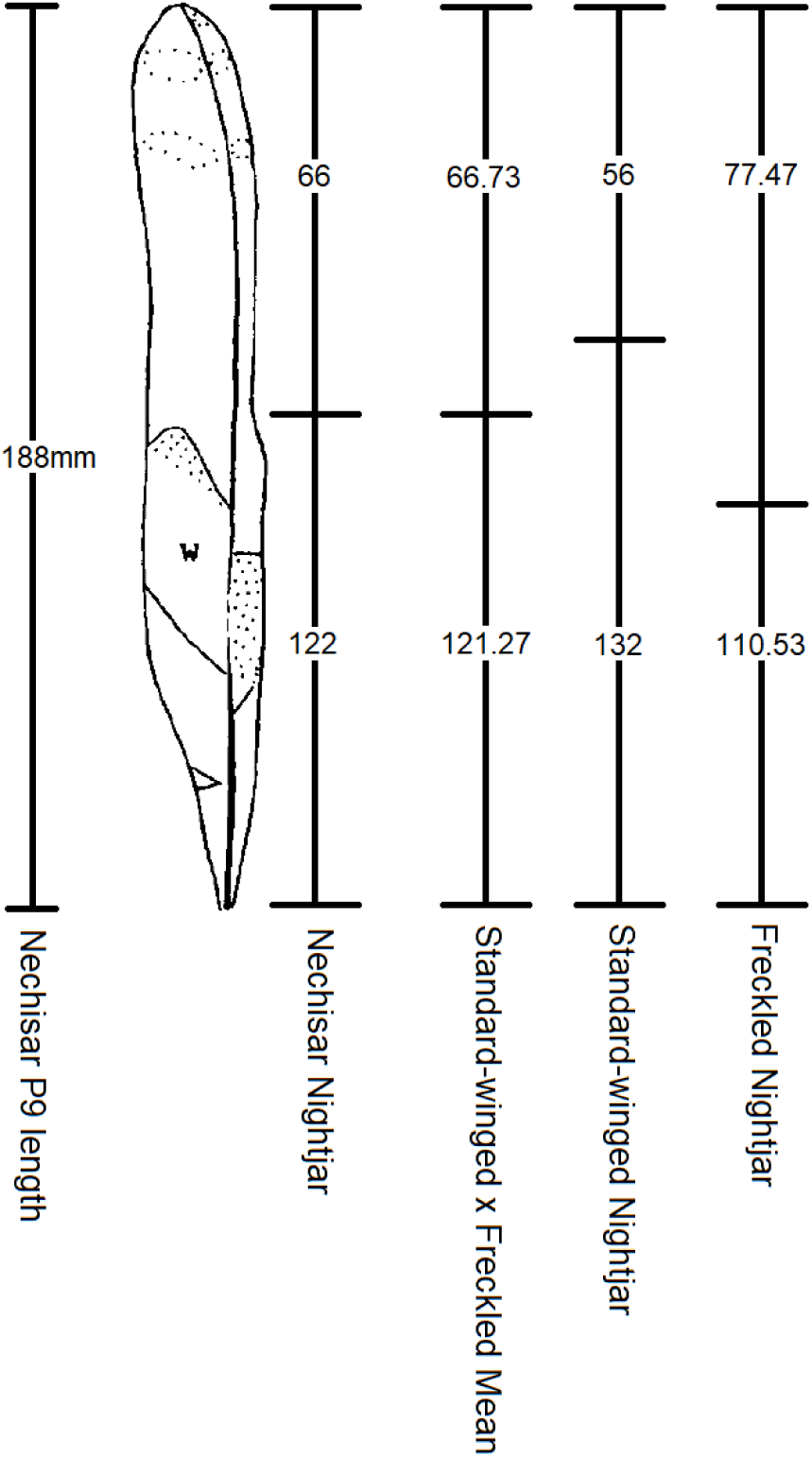
The relative position of the emargination on the P9 feather of an equal length to that of the Nechisar holotype. Shown are the emargination position of the Nechisar holotype, the mean emargination positions of Standard-winged Nightjar and Freckled Nightjar, and finally an average between the ratios of those two parent taxa. The Nechisar holotype measurements are shown to be nearly identical to the mean position between the two putative parent taxa.

The cryptic plumages of nightjars can superficially appear very similar, but both Freckled and Standard-winged Nightjars exhibit distinctive plumage patterns that are recapitulated in Nechisar Nightjar. The primary feathers of Standard-winged Nightjars are generally uniformly ‘tiger-striped’ cinnamon-ginger and blackish-brown, with colder dark-grey vermiculation patterns replacing the cinnamon-ginger on the distal tips of outer primaries, generally most prominent on P9-P5 (Figure 3). There are no white primary patches in Standard-winged Nightjars. In Freckled Nightjars, strong ‘tiger-striping’ is restricted to the inner primaries. All primaries have distal areas of colder vermiculation, but in Freckled Nightjar the outer four primaries are largely blackish brown with, in males, white patches variable suffused orange/brown (Figure 3). In the Nechisar Nightjar the primary patterns resemble very closely that of Freckled Nightjar – the outer four primaries are largely blackish-brown, with the four white-orange buff white spots, and barring and vermiculation restricted to the distal tips. The inner five primaries are variable tiger-striped dark brown and cinnamon-ginger, with cold dark grey vermiculation replacing the cinnamon bars at the primary tips. The plumage pattern is entirely consistent with Freckled Nightjar being one parent of the Nechisar Nightjar, and contrasts for example other potential parents such as European Nightjar, where the tiger-stripe patterns persist on the outer primaries 6-10 (Figure 3).

## Discussion

### Hybrid origin of Nechisar Nightjar

Based on the genetic evidence, Nechisar Nightjar holotype, a single wing, represents a hybrid between a female Standard-winged Nightjar, and another *Caprimulgus* species. The morphological and range-based study highlighted a single plausible candidate taxon for the paternal line, the Freckled Nightjar. This was also one of the 4 species for which no nuclear data was available.

The cytb sequence of Nechisar Nightjar was not 100% identical to any previously sequenced Standard-winged Nightjar, but this is most likely because Standard-winged Nightjars are themselves very poorly sampled across their range and the geographic range of genetic variation within the species has not been mapped. There would remain a remote possibility that the Nechisar Nightjar is a valid taxon, that is very closely related to the Standard-winged Nightjar, such that it either has ongoing gene flow or has only recently speciated. The second haplotype for each of the nuclear markers were not particularly similar to known Standard-winged references, but the species is poorly sampled and may have a complicated lineage history, which could account for this variation. However, given the lack of any further sightings in any of the numerous subsequent expeditions to the Nechisar National Park, it would seem much more likely that this unique specimen was exactly that, a one-off hybrid. It is an exceptional discovery - indeed this would, we believe, represent the first confirmed hybridisation event within Old World nightjars. However, given the nocturnal and cryptic life history and morphology of nightjars, it is possible that hybridisation events occur with some regularity, but have avoided detection.

In the wider Strisores, hybridisation is well documented, particularly in hummingbirds (e.g., Banks & Johnson 1961; Donegan & Dávalos 2011; Graves et al. 2016; van Dort & Juarez-Jovel 2016), for example as described for the Bogota Sunangel above.

The Nechisar Nightjar holotype represents the second single specimen *Caprimulgus* species to be shown through genetic techniques to be invalid in recent times, after Vaurie’s Nightjar (Schweizer et al. 2020). This highlights the utility of molecular methods as an aid for traditional taxonomy, and in no way detracts from the large amount of work performed for the description of the specimen in Safford et al. (1995). The resolution of invalid species designations is vital from a conservation perspective, particularly as resources are extremely limited and need to be correctly focused on preserving true biological and genetic diversity. The discovery of the Nechisar Nightjar is a once-in-a-lifetime experience and great credit must go to its finders for realising what a unique specimen they had. In the 1991 expedition, finding a new nightjar species in the Nechisar National Park was probably more likely than finding a previously unknown hybrid. The biology of a male Freckled Nightjar, lacking ‘standards’, successfully mating with a female Standard-winged Nightjar, raises a lot of questions, but it seems very likely that this has happened, at least once. Similar examples of unlikely hybridisation between taxa that otherwise exhibit very high levels of discrimination and sexual selection have been well described in birds of paradise (Paradisaeidae) (Stresemann, 1931; Martin (2015; Blom, et al., 2024; Thörn et al., 2024).

### Taxonomic implications of the mitochondrial phylogeny

The mitochondrial phylogenies produced as part of this study represent the most complete coverage of European and Afrotropical nightjar species to date, and revealed some interesting results. Firstly, the deep divergence within European Nightjar mirrors the findings of Schweizer et al. (2020) and may indicate that European Nightjar may in fact consist of two cryptic species, a sandy and pallid Eastern population (‘Vaurie’s), and a darker and more contrastingly marked Western population. The placement of the European Nightjar clade as sister to Plain Nightjar *C. inornatus* was also a novel finding and differs from previous phylogenies. Han et al. (2010) placed European Nightjar as sister to Rufous-cheeked Nightjar *C. rufigena*, but it appears as though the European Nightjar sample used in that study had been misidentified, as it had cytb sequence which was divergent from all other European Nightjar sequences both on Genbank and produced in this study.

A similar pattern was detected within Nubian Nightjar, with a pale and sandy migratory population in the Levant and Sudan separated from a darker plumaged sedentary population in East Africa and Socotra. The case to treat these two populations as separate species is enhanced by the paraphyly caused by the surprising placement of Star-spotted Nightjar as sister to the East African Nubian population. Given this unexpected pattern and the fact that the phylogeny was constructed based on a single mitochondrial gene means that the findings should be treated with caution and a more comprehensive phylogenetic study of this clade is warranted before any taxonomic decisions are made.

Two complexes within this study have been the subject of ongoing taxonomic debate. The Slender-tailed/Square-tailed/Long-tailed Nightjar complex has historically been treated as one, two and three species. The findings of our molecular study confirm the distinctiveness of the Long-tailed Nightjar, while the level of divergence between Square-tailed and Slender-tailed Nightjars was very low, and this may suggest that these two taxa may in fact be conspecific.

The Fiery-necked/Montane/Black-shouldered Nightjar complex has also been the subject of conflicting taxonomic assessments. The Rwenzori Nightjar *C. poliocephalus ruwenzorii* has been considered a species in its own right and a subspecies of Montane Nightjar. This study would suggest that Rwenzori Nightjar is a distinct species, while the taxonomy of the Fiery-necked, Black-shouldered and Montane Nightjar is much more ambiguous, with no clear structure visible in the clade in either of the produced phylogenies. The taxonomy of this group is still in flux, as evidenced by the lumping of Black-shouldered Nightjar and Fiery-necked Nightjar by IOC at the start of Nov 2022 (Gill et al. 2022). A more comprehensive phylogeographical study, targeting a larger number of loci is required to resolve this complex more conclusively.

### Limitations and future work

We continue to attempt to source material from Freckled Nightjar, and this preprint represents a cry for help in locating a fresh sample. The mitochondrial study forms the basis of a full phylogeny of Old World nightjars. However, given the cryptic nature of the entire group, wider genomic coverage of all subspecies from all across each species range would yield a much more robust phylogeny and would be likely to highlight a number of deep splits which may be cases of cryptic speciation. Within the limited dataset collected in this study, deep splits have been revealed in Nubian Nightjar and European Nightjar, and the placement of Star-spotted Nightjar within the Nubian Nightjar clade was a particular surprise. Further genomic study is required to confirm the legitimacy of this evolutionary relationship.

## Acknowledgements

This work was performed with funding to MC and TJS from The Sound Approach PhD Studentship, whose support is gratefully acknowledged. We thank multiple people from academic, museum and ornithological backgrounds who have supplied information and specimens.

## Supplementary Material

**Supplementary Figure 1.**
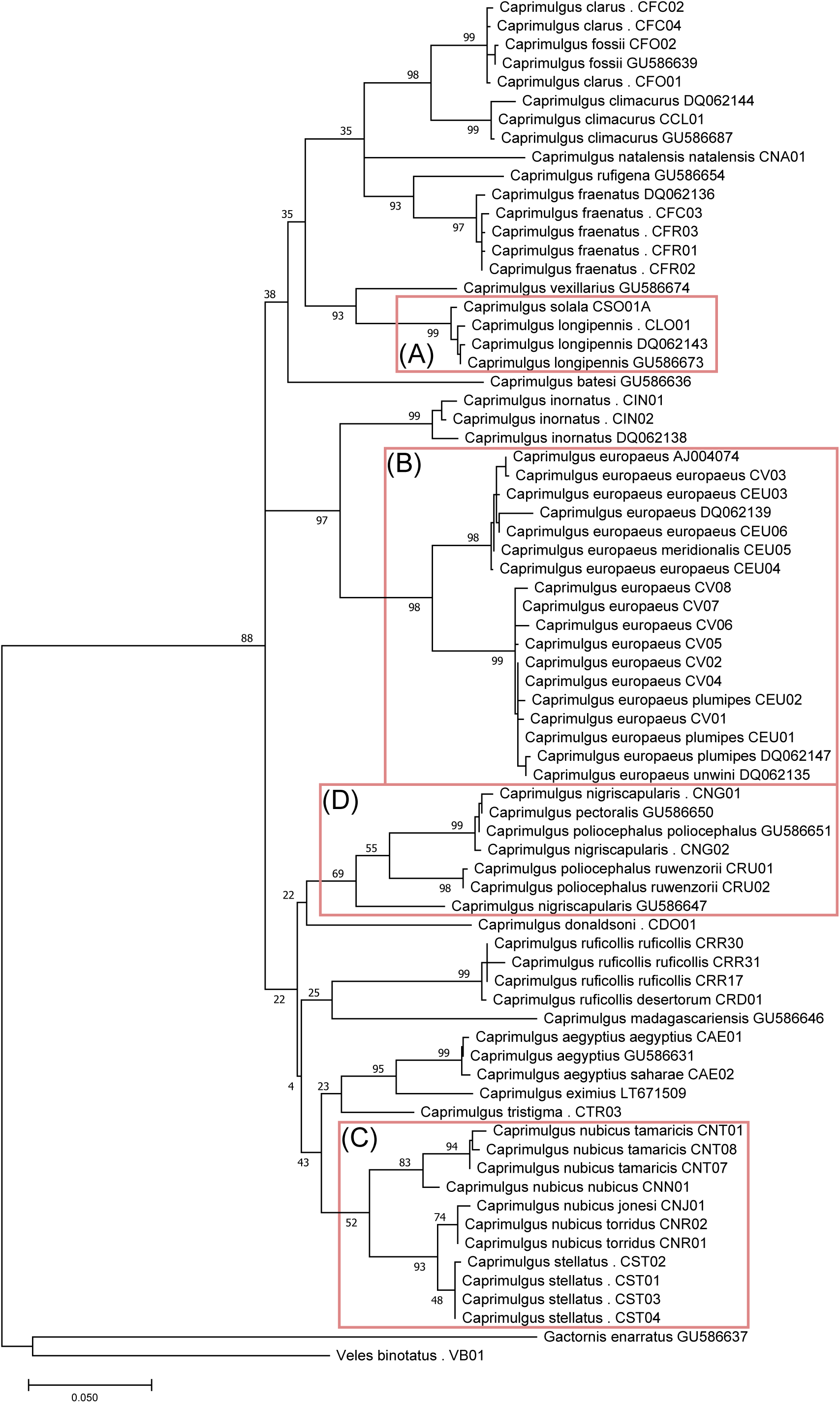
Maximum Likelihood phylogeny of Afrotropical and Eurasian nightjars constructed using MEGA X, based on the barcoding region of the mitochondrial cytochrome b gene. Bootstrap values are presented at the nodes. Clades of note are highlighted in red boxes: (A) Nechisar Nightjar is, at least maternally, a Standard-winged Nightjar; (B) deep divergence observed within European Nightjar; (C) paraphyly in Nubian Nightjar complex; (D) poor resolution in the Black-shouldered/Fiery-necked/Montane Nightjar complex.

**Nechisar Nightjar Cytb Sequence:**

CTGATGAAACTTCGGATCCCTCCTAGGAATCTGCTTGGCAACACAAATCCTAACCGGACTCCTCCTAGCCACACACTACACCGCAGACACAACCCTTGCCTTCTCATCCGTCGCCCACACCTGCCGTAATGTTCAATACGGCTGACTAATTCGCAACTTACACGCAAACGGAGCATCATTTTTCTTCATTTGCATTTACCTTCACATTGGACGAGGCCTATACTACGGATCATACCTTTACAAAGAAACCTGAAACACAGGAGTCATCCTCCTACTCACCTTAATAGCAACTGCCTTCGTAGGCTACGTCCTACCATGAGGACAAATATCATTCTGAGGAGCTACAGTCATCACCAACCTATTCTCAGCCATCCCATACATCGGCCAAACCCTTGTAGAATGAGCATGAGGTGGATTTTCTGTAGACAACCCTACACTAACCCGATTCTTCGCCCTACACTTTCTACTCCCCTTTATAATCGCCGGCCTTACCCTCATCCACTTAACATTCCTTCACGAATCAGGCTCAAACAACCCCCTGGGAATCGTATCAAACTGCGACAAAATCCCATTCCACCCTTACTTCTCCCTAAAAGACATTCTAGGCTTCGCACTAATACTTCTCCCATTAATAGCGCTCGCCATATTTTCCCCAAACCTACTAGGAGACCCAGAAAATTTCACCCCAGCAAACCCACTAGTCACACCCCCACATATTAAGCCCGAGTGATACTTCCTGTTTGCATATGCTATCCTACGCTCAATTCCAAACAAACTAGGAGGAGTCTTGGCCCTCGCTGCCTCCGTCCTAATTCTCCTACTAATCCCCCTCCTACACAAATCCAAACAACGCACACTAACTTTCCGCCCCCTCTCCCAACTACTATTCTGAACCCTAGTTGCCAACCTAATTATCCTAACCTGAGTGGGCAGCCAACCAGTAGAACACCCATTCATTATCATTGGC CAACTAGCCTCCCTCACCTACTTCTCCATCCTCCTAATCTTATTCCCCACCATCGGGGCCCTCG

